# Optimizing scheduling in dual-pulse nucleoside labeling experiments for cell cycle analysis

**DOI:** 10.1101/2025.07.14.664660

**Authors:** Alastar Phelan, Constandina Pospori, Cristina Lo Celso, Chiu Fan Lee

**Affiliations:** Department of Bioengineering, Imperial College London, South Kensington Campus, London SW7 2AZ, U.K; Department of Life Sciences, Imperial College London,South Kensington Campus, London SW7 2AZ, U.K

## Abstract

All eukaryotic cells go though a universal sequence of phases during their division cycle, where the phase timings vary according to cell type and state. Dual-pulse nucleoside labeling (DPNL) is a standard, widely applicable experimental technique to probe cell cycle kinetics at the population level, including in living organisms. In such an experimental protocol, a key scheduling parameter is the choice of waiting time between the two labeling pulses. Here, we use simulation to show that the waiting time can be optimized with regards to the signal-to-noise ratio of the cycle parameters as inferred from experimental outputs, which is of particular importance in DPNL experiments with small numbers of cells and repeats. We further discuss the procedure to perform such a task in an experimentally relevant setting.

---

Tightly regulated cell divisions are crucial to the development and maintenance of all organisms. Before a cell can divide, it must go through multiple phases with carefully controlled checkpoints – termed the G_1_ (1st gap phase), S (the synthesis phase when DNA is replicated), G_2_ (2nd gap phase), and M (for mitosis). A quantitative understanding of how much time a cell spends in these distinct phases is fundamental to our understanding of basic cellular functions, from DNA replication to organelle duplication.

Given the importance of understanding cell cycle dynamics, diverse experimental techniques have been developed to quantify the process [1], among these the use of nucleoside substitution labeling is particularly popular due to its ease of use and high sensitivity [2]. In such an experimental procedure, cells replicating their DNA in the S phase will incorporate modified nucleosides (e.g. EdU (5-Ethynyl-2’-deoxyuridine), a thymidine analog, Fig. 1a) whose presence can then later be detected (e.g., via flow cytometry methods). Detecting the amount of cells with these modified nucleosides in an EdU pulse-chase experiment has been a gold standard way to quantify the duration they spend in the S phase [3].

**FIG. 1.**
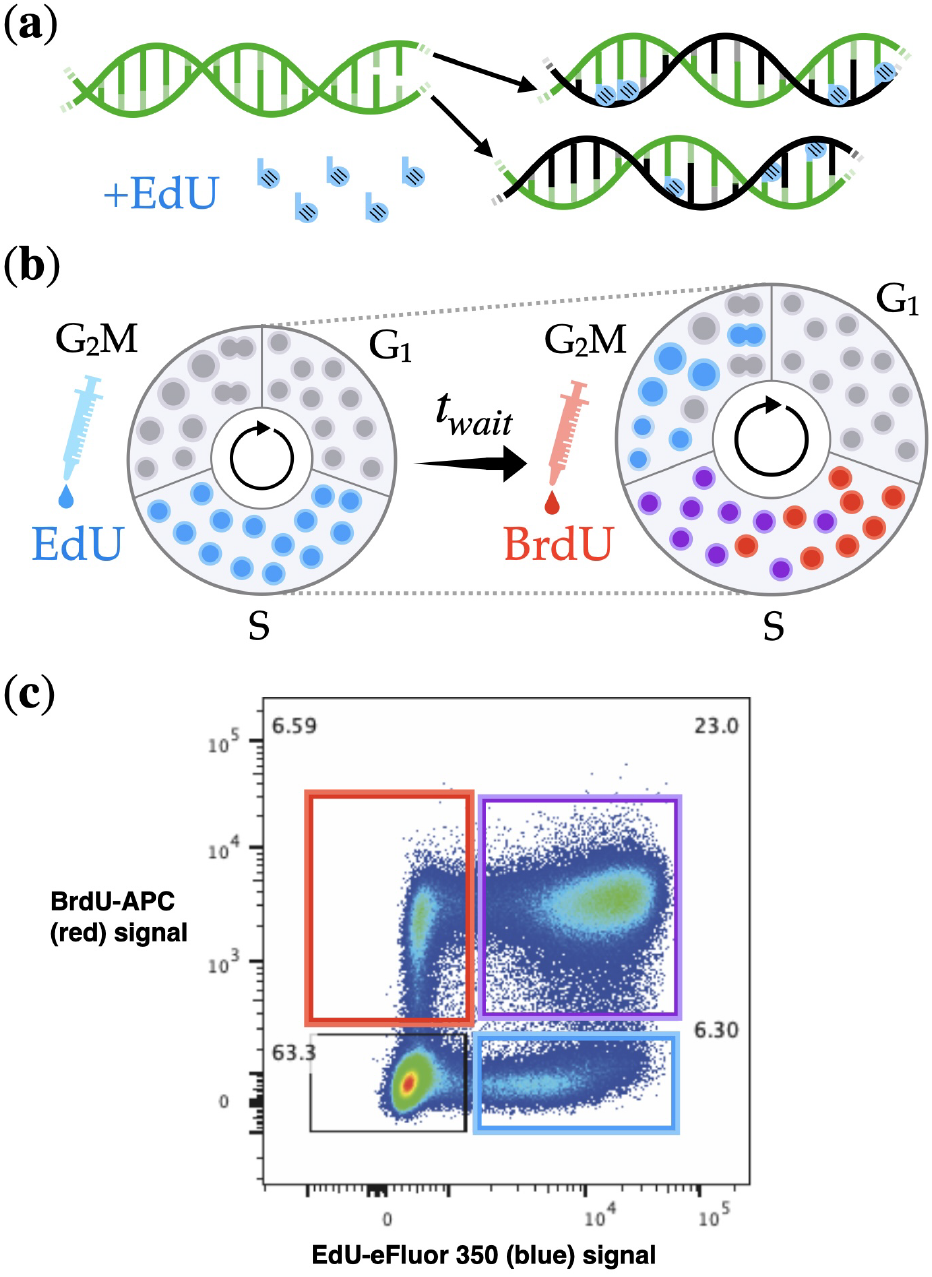
Nucleoside analog incorporation in Dual Pulse Nucleoside Labeling (DPNL) to study the cell cycle. **(a)** If thymidine analog EdU (5-Ethynyl-2’-deoxyuridine) is available during DNA synthesis, it is incorporated into each new half-strand produced. **(b)** In DPNL, the progress of cells through the cell cycle can be tracked by using a pulse of EdU (blue) followed by BrdU (5-Bromo-2’deoxyuridine, red) to show which cells have finished (blue), remained in (purple), or newly begun (red), DNA synthesis in the time between pulses. **(c)** The typical output of a dual pulse labeling assay is a square-shaped distribution of fluorescence intensity, as measured by flow cytometry, which is proportional to the number of analog molecules. There are four density peaks at the coarsest level [7, 8] corresponding to the colorings shown in Fig. 1b. The percentage of cells with each fluorescence combination are shown in text near to each box.

To directly probe cell cycle dynamics, a sequential substitution of nucleosides using two distinct labels (e.g., EdU and BrdU (5-Bromo-2’-deoxyuridine)), termed Dual Pulse Nucleoside Labeling (DPNL) has been developed [4, 5]. In this experimental procedure, the second label is introduced into the system at a time *t*_*wait*_ after the introduction of the first label (Fig. 1b). As a result, four distinctly labeled cell groups (see Fig. 1c) can be detected (as opposed to two in the single labeling method), boosting the information output of the experiment and especially improving S phase inference accuracy [6]. Doublepositive cells (purple, Fig. 1b & c) are those which were in S phase during both EdU and BrdU exposure; single-positive (blue, red in Fig. 1b & c) are those which were only in S phase at either the EdU or the BrdU exposure time, meaning they were late S phase or early S phase respectively at the corresponding pulse time; otherwise cells are double-negative. For populations engaged in a synchronized cell cycle [9], only one of these four label states would be observed in a DPNL assay. While BrdU-positivity at the end of a DPNL experiment implies that a cell is in S phase, both the EdU-single-positive and the double-negative populations are split between the G_2_M and G_1_ phases in continuously proliferating cell types. This output structure suggests that S phase inference will be better constrained by DPNL data compared to G_1_ and G_2_M.

However, cells in vivo are rarely synchronized, and variability (even purely at the population level) in the times that cells within a population take to complete each phase, due to constraints such as tissue crowding [10, 11] or mitogenic factors [12, 13], remains a challenge for quantitative modeling where detailed timeseries data are scarce [14]. Noisy cycle dynamics can have a large impact on inference about a whole population when the number of specific cells of interest (e.g. actively proliferating haematopoietic stem cells in a mouse) may be fewer than 1000 cells [11]. Compounding with this, the number of repeats of cells or tissues sampled is typically small in a given study, highlighting the need to optimize the accuracy of the experimental measurements. Indeed, a clear control parameter in this dual-labeling method is the waiting time *t*_*wait*_ between the introductions of the two labels. A longer *t*_*wait*_ allows more EdU-positive cells to leave S phase before the BrdU pulse, giving a smaller doublepositive population, but larger single-positive populations, for *t*_*wait*_ shorter than S phase. However, how to choose *t*_*wait*_ to optimize the information output has thus far not been thoroughly investigated. Here, we perform this task and demonstrate, using simulation of a simple model of cell cycle (Fig. 2), how to find the *t*_*wait*_ that optimizes the signal-to-noise ratio (SNR) of the experimental measurements. Our work provides a proof of principle in how to use modeling to improve the precision of a widely used experimental method in the study of cell cycle dynamics.

**FIG. 2.**
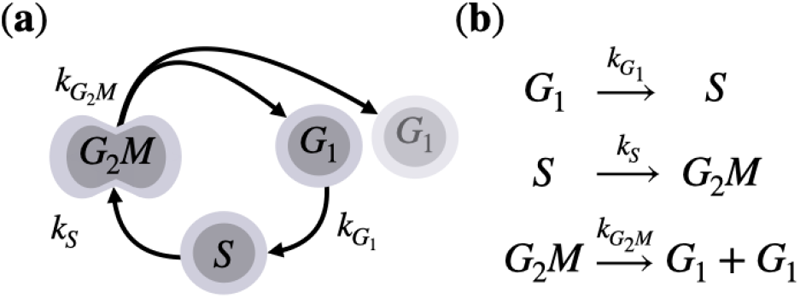
The three species, three parameter model of the cell cycle we have used in this work. **(a)** Cells in each stage *i* of the cycle advance to the next at a rate *k*_*i*_, as a Poisson process. Two new cells are generated at the end of the cycle, both of which continue to advance through the cycle phases from the beginning, G_1_ phase. **(b)** Reaction scheme describing the cell cycle phase transitions. As a 3-stage Poisson process, the total duration of the cycle is Erlang-distributed [18], while using the minimum number of parameters to keep results fully interpretable.

## I. MODELING THE CELL CYCLE DURING A DUAL-LABELING EXPERIMENT

We model the progress of the cell cycle through distinct phases equivalently to chemical reactions with rates *k*_*i*_ (Fig. 2), subject to fluctuations in the reaction kinetics (see details in Appendices A & B). We have used a common simplification in the model, namely contracting G_2_ and M phases into a single phase, termed G_2_M. The two phases, usually the shortest in the cycle, are not readily distinguishable when looking solely at EdU fluorescence or DNA content [1, 15], hence the phases are often modeled as one.

Cells are modeled as progressing through the cycle asynchronously, which is accepted for quantifying the cell cycle across a cell population in most cases [1, 2, 9]. The model’s strength is in its simplicity, where any other noisy cell cycle model is necessarily more complex [16, 17]. Pure DPNL’s output has two channels, giving four measurable quantities: the total number of cells with each fluorescence combination (Fig. 1 b & c), meaning any attempt to infer more than four independent model parameters from the output is likely to be difficult.

To find the model parameter-dependent statistics of the labeled cell counts at the end of DPNL experiments, so that we can investigate how noise can be mitigated in the process, we model DPNL at the population level stochastically. The Master equation describing probabilistic trajectories of the population state, described by a vector 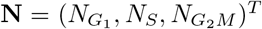, the number of cells in G_1_, S and G_2_M phase respectively, is as follows:

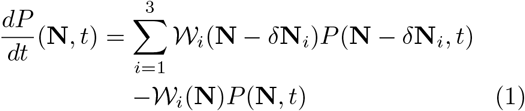

where *P* (**N**, *t*) is the probability at time *t* of having a population state **N**. *𝒲*_*i*_(**N**) = *k*_*i*_*N*_*i*_ is the transition rate out of the state **N** via a single cell transition *δ***N**_*i*_, the loss of one cell and the gain of one or two in the next phase as described in the reaction scheme Fig. 2b, with *N*_*i*=1_ corresponding to the number of G_1_ phase cells, *k*_*i*=1_ being their rate of progression to S phase and so on for *i* = 2, 3.

For the labeling part of the simulation, we apply an EdU label to all cells in the S phase at time *t* = 0 only and apply BrdU to all cells which are in S phase and either EdU positive or negative at *t* = *t*_*wait*_ only. In other words, labeling is assumed to be instantaneous and complete, as supported experimentally [11]. We do not account for EdU dilution after cell division since when analyzing flow cytometry data, the threshold for EdU-positivity can be chosen while taking the dilution into account, with even the most diluted cells readily distinguished from EdU-negative cells. All labeled cells continue to advance through the cell cycle at unaltered rates until the end of the simulation, ultimately generating a separately cycling population for each label combination by this time. Given the chemical master equations and labeling scheme above, we will now use the Gillespie algorithm [19] (see Appendix B for details) to simulate an array of dual pulse nucleoside labeling experiments.

## II. GENERATING A LOOK-UP TABLE FROM KINETIC RATES TO DPNL OUTPUTS

As we focus here on population noise, which manifests itself most strongly in the study of small cell colonies, e.g. actively proliferating stem cells, we initialize our system with 300 cells, consistent with the reported numbers of haematopoietic stem cells entering S phase per hour in the mouse hind leg [11], as an example of a small population of replicating cells. The initial composition corresponds to the number of cells in each phase in the steady growth solution from solving for the average dynamics of the Master equation (details in Appendix C). Further motivated by the typical experimental procedure [11, 20, 21], we have a 30-minute interval after BrdU administration before counting up the cells with each label combination, which ensured the complete incorporation of BrdU in real experiments.

The total number of cells with each of the four label combinations (EdU single-positive (blue), BrdU single-positive (red), double-positive (purple), double-negative (gray)), is the output of an experiment (see Fig. 1b & c), which can be used to calculate individual phase durations, with the exact calculation depending on the cell type [4, 16, 20, 22, 23].

To probe the statistics of labeled cell outputs, we simulate across a range of scenarios by sweeping across a grid of model parameters *k*_*i*_ and *t*_*wait*_, while fixing the overall cell cycle length to 24 hours. The parameter sweep range considered (details in Appendix D) is within the experimentally relevant range based on observed ratios between G_1_, S and G_2_M phase duration [24, Table S1] and keeping *t*_*wait*_ ≤ 12 hours. Beyond this limit, a significant number of cells could both exit S phase after receiving the EdU pulse and re-enter it in another lap of the cycle by the time of the BrdU pulse. These cells would register as double-positive and could make cell count outputs difficult to distinguish from a slower case where the same number of cells have remained in S phase throughout to register as double-positive.

For each combination of model parameters, *k*_*i*_ and control parameter *t*_*wait*_, we generate its probability distribution of output cell numbers through 10,000 repeats. This dataset generated will serve as a ground truth look-up table when we deal with typical experimental studies where the small number of repeats adds another layer of noise to our consideration.

## III. INFERRING CELL CYCLE TRANSITION RATES FROM TYPICAL EXPERIMENTS

We now use the ground truth dataset generated as a dictionary to enable us to perform the inference of the model parameters *k*_*i*_ from the outputs of a typical experiment, which consists of only around 3-5 repeats. Hence, we perform 3-repeat simulations for each model parameter combination. For every 3repeat trial, we look up the closest entry (see Appendix D for details) in the dictionary data previously collected to infer the most likely *k*_*i*_. To obtain the statistics of this inference process, we perform a large number (*∼* 10^3^) of 3-repeat trials of simulations per model parameter combination to enable us to build up a distribution of estimated *k*_*i*_ from the simulated 3-repeat trials. Investigating these distributions will enable us to decipher the SNR of our inference process as a function of the waiting time *t*_*wait*_.

## IV.PICKING WAIT TIME TO OPTIMIZE THE SIGNAL-TO-NOISE RATIO

In a typical experiment, since the number of repeats is small, the inferred parameters *k*_*i*_ will deviate from the true cell cycle rates due to intrinsic fluctuations.

Consider counting the number of cells in a population before and after a growth period, which is controlled by the experimenter, in order to infer the population’s growth rate per cell. Waiting as long as possible is clearly the ideal case as the expected deviation of the average division time of a cell is suppressed with longer growth times in line with the central limit theorem. However, in DPNL, the cell cycle rate parameter inference cannot keep improving for longer waiting times because relatively fast-cycling EdU-positive cells re-entering S phase for the BrdU pulse can be difficult to distinguish from slow-cycling cells that remained in S during that interval, increasing error and sensitivity in the inference. Our simulation approach can quantify this trade-off.

To understand the impact of the stochasticity, we can use the inference SNR:

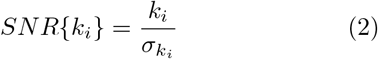

which corresponds to the ratio of the average most-likely *k* values and their standard deviations. Here, we can quantify the SNRs accurately by comparing results from our 3-repeat in silico experiment with the known ground truth rate parameters in our lookup.

In Fig. 3, we show the SNRs as a function of the waiting time *t*_*wait*_ for distinct set values of S phase and *G*_1_ phase periods, *t*_*S*_ (main plot) and 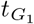 (inset), respectively. We find that *for any* set of initial ground truth parameters (color bar in Fig. 3), *SNR {k*_*S*_*}* is generically non-monotonic with respect to *t*_*wait*_, namely, there is indeed an optimal *SNR{k*_*S*_*}* at a nontrivial value of *t*_*wait*_. Although *SNR* 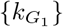 seems to show opposite trends to *SNR{k*_*S*_*}*, the local minima in *SNR* 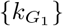do not coincide with the local maxima of *SNR k*_*S*_, thus indicating that there exists generally an optimal *t*_*wait*_ that can optimize the *SNR* of the inference process, depending on the experimental focus (e.g., whether it is on *t*_*S*_ or on 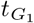).

**FIG. 3.**
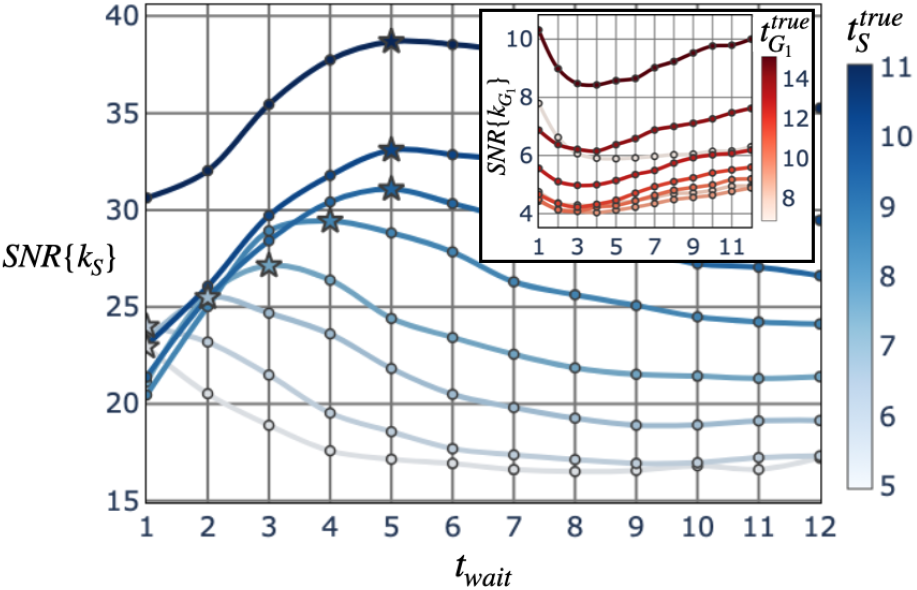
Signal-to-Noise Ratio of the lookup inference process for model parameters k_i_. **Main panel:** *SNR {k*_*S*_*}* vs *t*_*wait*_ for different S phase durations. A nontrivial S-phasetime-dependent optimum exists, reaching higher peak values for longer S phase durations, with the optimal pulse interval simultaneously becoming longer. **Inset:** Intermediate pulse interval times are unfavorable for G_1_ phase duration inference and hint at an indirect trade-off with S phase inference. The relative mean squared errors for each error are shown in Appendix E.

We will now describe how such an optimization can be applied in a typical experiment.

## V.A PROTOCOL IN OPTIMIZING THE WAITING TIME

The optimal waiting time between pulses in a DPNL investigation clearly depends on the cell cycle phase times of the cell types being assayed. One therefore would need to first start with an initial assumption of these times based on, e.g., existing estimates.

The next step is to agree on a quantity to optimize for (e.g., to focus on SNR of S phase inference or balance the strength of all parameters’ inference). Then the simulation procedure used in this paper can be used to obtain the optimal choice of *t*_*wait*_. The experimenter could run their experimental repeats using this choice.

One could of course also further optimize the whole procedure by repeating the procedure described in an iterative manner with input from experiments.

## VI. SUMMARY & OUTLOOK

By modeling the cell cycle as a sequence of three transitions with noisy timing, we have established a proof of principle that the waiting time between labels used in Dual Pulse Nucleoside Labeling can be nontrivially optimized for maximal Signal-to-Noise ratio of cell cycle kinetic parameter inference.

Future work could build on this result in multiple directions. 1) Performing a similar analysis, this time lifting the 24-hour cycle constraint, could have closer relevance to a wider host of cell types. 2) With a model that additionally quantifies nuclear DNA content via DAPI [25] at the end of a DPNL assay, the technique’s relative weakness for precise G_1_ and G_2_M phase inference is potentially resolved. 3) In addition, multi-parameter inference for a more complex model of the cell cycle has a higher chance of success with the DAPI-complemented DPNL data [16, 17]. 4) Lastly, incorporating differentiation to build a more interrelated picture of a cell replication system could help to ascertain the optimal dual labeling setup when the cell types being measured in a multiplexed flow cytometry setup have different cell cycle kinetic parameters to be inferred [26].

## ACKNOWLEDGMENTS

We have used the Imperial College Research Computing Service for this project (DOI: 10.14469/hpc/2232). For funding support, DP, CLC, & CFL thank CRUK Imperial Centre & NIHR Imperial BRC Data Science in Cancer Research Award 2020; DP thanks CRUK Development grant; CLC thanks Wellcome Investigator award [212304/Z/18/Z] and CRUK Programme foundation award [C36195/A26770].

## APPENDIX A: SOLVING THE FULL MASTER EQUATION

In terms of probability current, where state **N**’s outgoing current 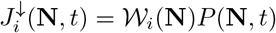,

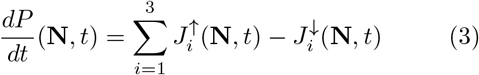

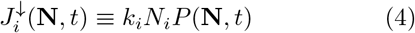

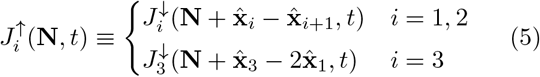

with *P* (**N**, *t*) the probability at time *t* to occupy a state described by vector 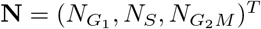, the total number of cells in each cycle phase; *k*_*i*_ the rate parameter for a cell advancing from phase *i* to 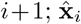 represents a single cell in phase *i*. The expectation of a quantity at a time *t* can be solved by multiplying the master equation by that quantity and summing over states. For example, the mean number of G_1_ phase cells at a time *t* can be calculated as

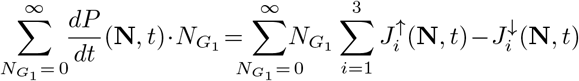

Since the number of G_1_ cells is just a quantity to be counted over, the time derivative and sum on the left-hand side can be commuted

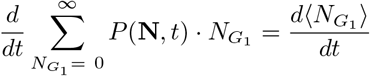

and expanding the probability current terms on the right hand side, with a change of index we find that the expression can be simplified

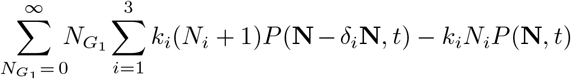

where *δ*_*i*_**N** represents the change of cell counts in each phase brought about by one cell making the phase advancement *i* as described in (5). The *i*^th^ element of *δ*_*i*_**N** is always *−*1. Making the substitution **N***− δ*_*i*_**N** *→***N**, noting that boundary cases where *N*_*i*_ = 0 for any *i* are handled naturally by the mass action form of the master equation, the expression becomes

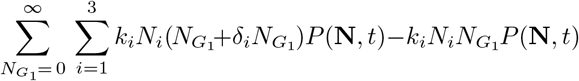

now the sums can be evaluated and a cancellation made

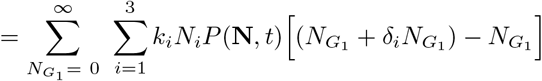

using that 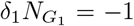 and 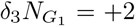, we recover the deterministic noise-free limit of the model equations,

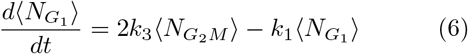

To calculate higher statistical moments, the same pro-cess is followed but with multiplying by 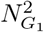 for ex-ample instead. As a linear system, the moments of the3-phase cell cycle model are summarized in equations (7) and (8) below.

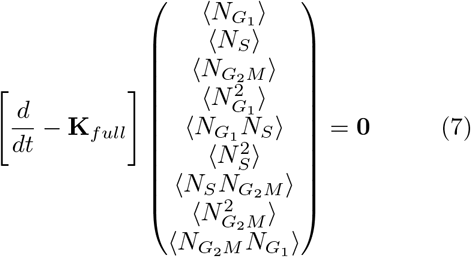

The resulting ODEs are solved by a numerical integration technique for convenience in producing the following plots, which show excellent agreement between Gillespie simulations and our analytic results from the approach outlined above, when quantifying the mean and standard deviation of the total number of cells with each label combination.

At the labeling times, the expected number of S phase cells *⟨ N*_*S*_*⟩* and expected squared number of S phase cells 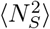 is transferred to the population state of the newly labeled population, with these set to zero in the source population i.e. colorless cells exposed to either label or EdU-positive cells exposed to BrdU in the late stages of an experiment. Cross-terms 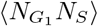 and 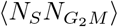 are set to zero. This reproduces the dynamics seen in simulations (Fig. refanalytic-plots).

Getting from model parameters to the average and standard deviation of the labeled cells at the end of a DPNL experiment stops one step short of the DPNL simulations, which noisily infer rate parameters from the labeled cells. Further work could look at this process analytically, to predict where the peak in signalto-noise ratio for any parameter inference should be.

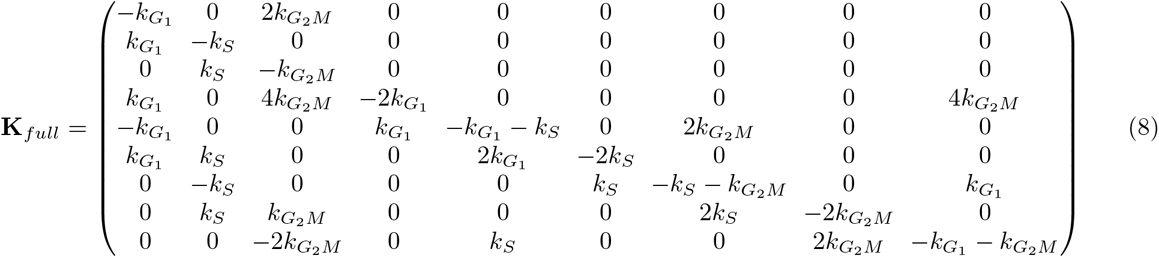

**FIG. 4.** Full rate matrix for the average dynamics of number of cells in each phase as well as their squares and crossproducts, which are used to calculate the expected variance at a given time. The columns and rows proceed in the order 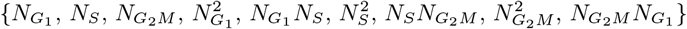

**FIG. 5.**
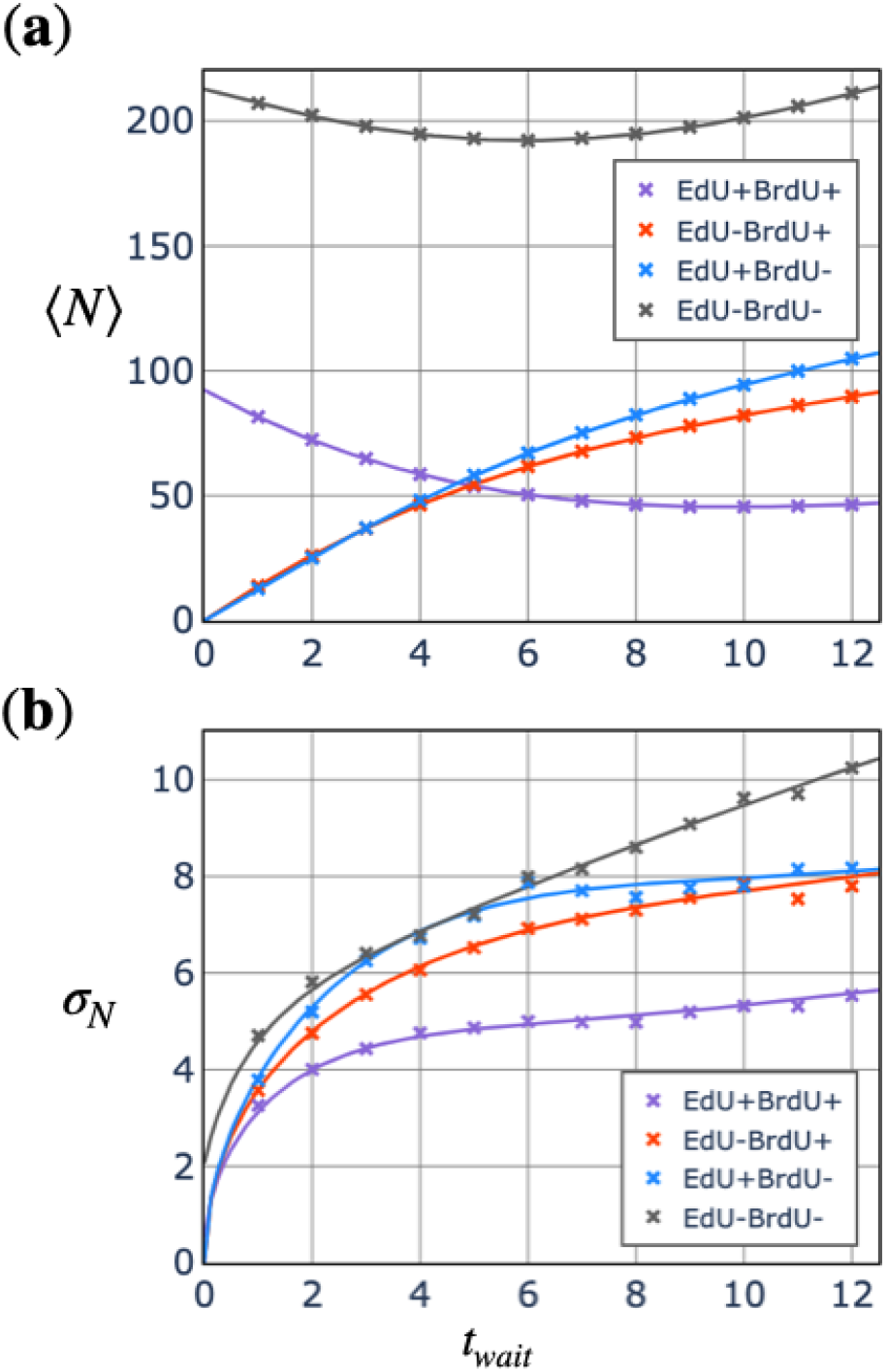
**(a)** Average and **(b)** standard deviation of cell counts for different inter-pulse waiting times from simulation (crosses) and the analytic solution (lines). The signal-to-noise ratio of 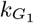 and *k*_*S*_ parameter inference is a non-trivial function of the quantities plotted above.

## APPENDIX B: STOCHASTIC SIMULATION ALGORITHM

The core Gillespie algorithm [19] for our simulated dual pulse nucleoside labeling experiments is as follows. At each time until the end of the experiment,

1. Calculate the current propensities (bulk rates) of the system, 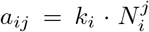, where *i* iterates over the reaction index (equal to the phase index), and *j* iterates over the four fluorescent label combinations i.e. cells which are positive for EdU, BrdU, both, or neither.
2. Draw a random number from an exponential distribution centered on (Σ_*i,j*_ *a*_*ij*_)^*−*1^, add it to the current time of the simulation.
3. Generate a random number between 0 and Σ_*i,j*_ *a*_*ij*_, find the first element in the cumulative sum which exceeds the random number in order to pick which single reaction takes place in this time interval, then add and take away the appropriate cells according to the stoichiometry of that reaction. For example, picking random number 5.0 with reaction propensities 1.0, 3.0, 2.0, 4.0 will result in the third reaction taking place.

The initial state of the system, the steady growth state for 300 cells with given model parameters, is exposed to EdU at time 0. This means making the assignments 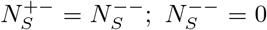 While the above population dynamics are running, at each time step, the following two conditional actions are performed to simulate the experimenter’s labeling and harvesting steps.

- Otherwise, if the BrdU labeling time was just passed for the first time, set 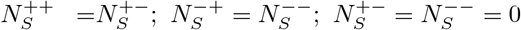.
- Otherwise, if the end time for the experiment, *t*_*wait*_ + 0.5 (cells cycle for a further half an hour for fixation as in real experiments), was just passed, then report the end state as the number of cells with each label combination: 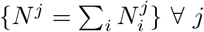. End the simulation.
- Otherwise, go to next iteration.

## APPENDIX C: STEADY GROWTH STATE

In contrast to a cyclic deterministic mass-conserving reaction scheme [27][28], the three stage cell cycle model has no steady state but instead can achieve balanced growth, which corresponds to the steady state of the normalized system

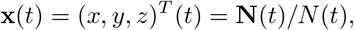

the number of cells in each phase of the cycle divided by the total number of cells at each time *t*.

With an exponential growth ansatz

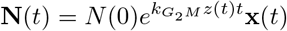

Differentiating with respect to time, the deterministic equations of motion from this ansatz become

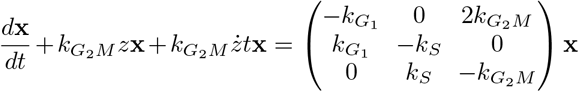

and the normalized system can be solved for a steady state 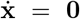 by *z → α*_1_ where *α*_1_ is the root of 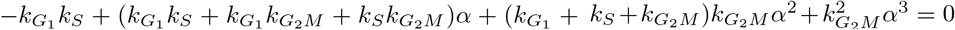 located near *α* = 1, and

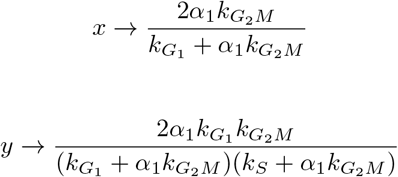

For a 24 hour cell cycle period with G_1_ lasting 11 hours, S 8 hours, G_2_M 5 hours [29], meaning 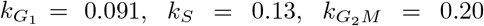, the renormalized steady state is situated at approximately *x →* 0.53, *y →* 0.31, *z →* 0.16. Note that G_1_ contains more cells than the other phases per hour of its duration. This is because the distribution of cells around the cycle at steady growth is biased towards earlier points in the cycle due to the generation of two new G_1_ cells at the end of G_2_M [1]. In the absence of any absorbing states since there is no cell death in our model, the stochastic trajectories remain close to the deterministic solution on average, within a thin tube of noise [30].

## APPENDIX D: PARAMETER SWEEPS

Parameters (*t*_*G*1_, *t*_*S*_, *t*_*G*2*M*_) were drawn from a hexagonal grid on the 2-simplex defined by *t*_*G*1_+*t*_*S*_ +*t*_*G*2*M*_ = 24h centered on (11.0, 8.0, 5.0) [31], resulting in 47, 35, and 13 unique values of each respective parameter, designed to cover a wide range of cell types with near 24 hour cycle periods. To convert to average rates 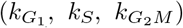, the reciprocal of each time is taken. 10^4^-fold repeat stochastic simulations build up a dictionary of most likely cell counts at the end of an experiment with each model parameter set

Running 3 repeats thereafter as if running a real resource-limited experiment and selecting the closest entry in the dictionary to the average cell counts across those 3 repeats, the stochastic behavior will lead to deviations from the real model parameters being selected as most likely. The scoring function to work out the closest parameter combination is

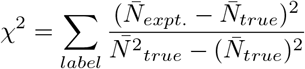

where subscripts denote whether the number comes from the 3-repeat experiment (expt.) or the large number of repeats (true). This process is repeated 1000 times per 3-repeat experiment per model parameter combination to find the mean and variance in inferred values of 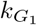 and *k*_*S*_. The ratio of the mean and standard deviation for each is then the inference signal-to-noise ratio for that model parameter.

APPENDIX E: INFERENCE ACCURACY

While the main focus of the paper is to optimize noisy cell cycle kinetic parameter estimation via a DPNL experimental output, we have also verified that any bias in the average inferred parameters is acceptably low. The mean error of the inference reduces sharply with longer *t*_*wait*_ as seen in (Fig. 6), with the worst-case scenario close to just 10% error for G_1_ phase duration inference for a waiting time of 1 hour. The worst-case scenario for S phase inference is a 2% error.

**FIG. 6.**
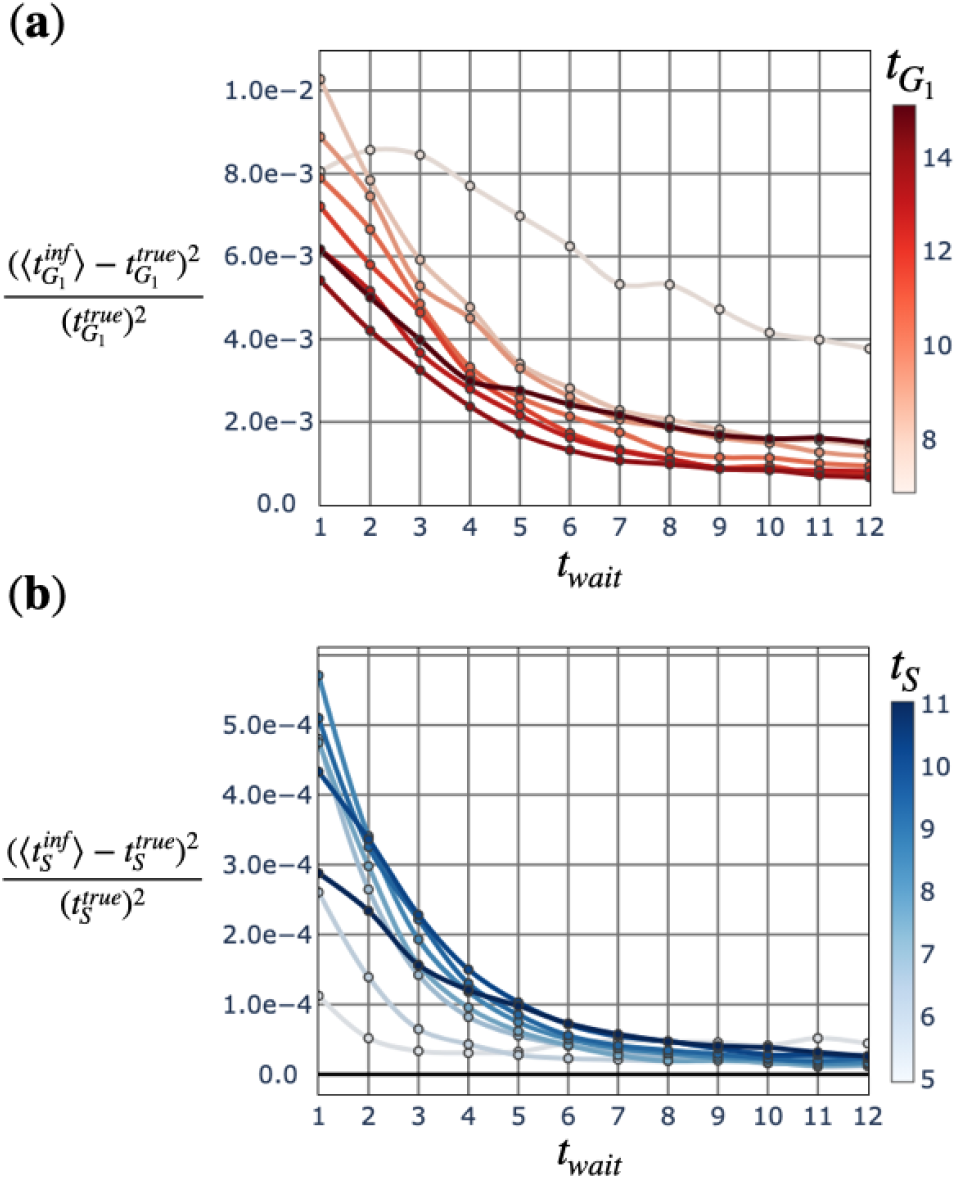
Squared mean error plots from parameter inference. **(a)** Normalized average error squared in G_1_ inference vs *t*_*wait*_, binned into 7 G_1_ duration groups. **(b)** Normalized average error squared in S inference vs *t*_*wait*_, binned into 7 S duration groups.

## Notes

### Competing Interest Statement

The authors have declared no competing interest.

## References

[1] Anna Ligasová, Ivo Frydrych, and Karel Koberna. Basic Methods of Cell Cycle Analysis. International Journal of Molecular Sciences, 24(4):3674, February 2023.

[2] Marta Bialic, Baraah Al Ahmad Nachar, Maria Koźlak, Vincent Coulton, and Etienne Schwob. Measuring S-Phase Duration from Asynchronous Cells Using Dual EdU-BrdU Pulse-Chase Labeling Flow Cytometry. Genes, 13(3):408, 2022.

[3] Leigh Mickelson-Young, Emily Wear, Patrick Mulvaney, Tae-Jin Lee, Eric S. Szymanski, George Allen, Linda Hanley-Bowdoin, and William Thompson. A flow cytometric method for estimating S-phase duration in plants. Journal of Experimental Botany, 67(21):6077–6087, November 2016.

[4] D. E. Wimber and H. Quastler. A 14C- and 3H-thymidine double labeling technique in the study of cell proliferation in Tradescantia root tips. Experimental Cell Research, 30(1):8–22, March 1963.

[5] Paolo Cappella, Fabio Gasparri, Maurizio Pulici, and Jürgen Moll. A novel method based on click chemistry, which overcomes limitations of cell cycle analysis by classical determination of BrdU incorporation, allowing multiplex antibody staining. Cytometry Part A, 73A:626–636, 2008.

[6] S. Kroll, D. Char, and S. Kaleta-Michaels. A stochastic model for dual label experiments: an analysis of the heterogeneity of S phase duration. Cell Proliferation, 28:545–567, 1995.

[7] Alexander D. Gitlin, Ziv Shulman, and Michel C. Nussenzweig. Clonal selection in the germinal center by regulated proliferation and hypermutation. Nature, 509(7502):637–640, May 2014.

[8] Oliver Bannard, Simon J. McGowan, Jonatan Ersching, Satoshi Ishido, Gabriel D. Victora, Jeoung-Sook Shin, and Jason G. Cyster. Ubiquitin-mediated fluctuations in MHC class II facilitate efficient germinal center B cell responses. Journal of Experimental Medicine, 213(6):993–1009, May 2016.

[9] Giansimone Perrino, Sara Napolitano, Francesca Galdi, Antonella La Regina, Davide Fiore, Teresa Giuliano, Mario di Bernado, and Diego di Bernado. Automatic synchronisation of the cell cycle in budding yeast through closed-loop feedback control. Nature Communications, 12:2452, April 2021.

[10] Carles Falcó, Daniel J. Cohen, JoséA. Carrillo, and Ruth Baker. Quantifying cell cycle regulation by tissue crowding. Biophysical Journal, 123:1–10, October 2024.

[11] O. Akinduro, T. S. Weber, H. Ang, M. L. R. Haltalli, N. Ruivo, D. Duarte, N. M. Rashidi, E. D. Hawkins, K. R. Duffy, and C. Lo Celso. Proliferation dynamics of acute myeloid leukaemia and haematopoietic pro-genitors competing for bone marrow space. Nature Communications, 9(1):519, February 2018.

[12] Ben Martynoga, Harris Morrison, David J. Price, and John O. Mason. Foxg1 is required for specification of ventral telencephalon and region-specific regulation of dorsal telencephalic precursor proliferation and apoptosis. Developmental Biology, 283:113–127, May 2005.

[13] Tim C. P. Somervaille and Michael L. Cleary. Identification and characterization of leukemia stem cells in murine MLL-AF9 acute myeloid leukemia. Cancer Cell, 10:257–268, October 2006.

[14] Chenhui Ma and Evren Gurkan-Cavusoglu. A comprehensive review of computational cell cycle models in guiding cancer treatment strategies. npj Systems Biology and Applications, 10:71, 2024.

[15] Ana Rita Araujo, Lendert Gelens, Rahuman SM Sheriff, and Silvia DM Santos. Positive Feedback Keeps Duration of Mitosis Temporally Insulated from Upstream Cell-Cycle Events. Molecular Cell, 64(2):362–375, October 2016.

[16] T. S. Weber, I. Jaehnert, C. Schichor, M. Or-Guil, and J. Carneiro. Quantifying the Length and Variance of the Eukaryotic Cell Cycle Phases by a Stochastic Model and Dual Nucleoside Pulse Labelling. PLOS Computational Biology, 10(7):e1003616, 2014.

[17] Adrien Jolly, Ann-Kathrin Fanti, Csilla Kongsaysak-Lengyel, Nina Claudino, Ines Gräßer, Nils B. Becker, and Thomas Höfer. CycleFlow simultaneously quantifies cell-cycle phase lengths and quiescence in vivo. Cell Reports Methods, 2(10):100315, October 2022.

[18] Christian A. Yates, Matthew J. Ford, and Richard L. Mort. A Multi-stage Representation of Cell Proliferation as a Markov Process. Bulletin of Mathematical Biology, 79:2905–2928, October 2017.

[19] Daniel T. Gillespie. Stochastic Simulation of Chemical Kinetics. Annual Review Physical Chemistry, 58:35–55, October 2006.

[20] Lachlan Harris, Oressia Zalucki, and Michael Piper. BrdU/EdU dual labeling to determine the cell-cycle dynamics of defined cellular subpopulations. Journal of Molecular Histology, 49:229–234, June 2018.

[21] Florian J. Weisel, Griselda Zuccarino-Catania, Maria Chikina, and Mark J. Shlomchik. A Temporal Switch in the Germinal Center Determines Differential Output of Memory B and Plasma Cells. Immunity, 44(1):116–130, January 2016.

[22] Mark A. Ritter, John F. Fowler, Youngmi J. Kim, Kennedy W. Gilchrist, Lawrence W. Morrissey, and Timothy J. Kinsella. Tumor Cell Kinetics Using Two Labels and Flow Cytometry. Cytometry, 16:49–58, 1994.

[23] Joaquín Martí-Cluá. Methods for Inferring Cell Cycle Parameters Using Thymidine Analogues. Biology (Basel), 12(6):885, 2023.

[24] Avraham Greenberg and Itamar Simon. S Phase Duration Is Determined by Local Rate and Global Organization of Replication. Biology (Basel), 11(5), May 2022.

[25] Vassilis Roukos, Gianluca Pegoraro, Ty C Voss, and Tom Misteli. Cell cycle staging of individual cells by fluorescence microscopy. Nature Protocols, 10:334–48, February 2015.

[26] Marta Rodríguez-Martínez, Stephanie A. Hills, John F. X. Diffley, and Jesper Q. Svejstrup. Multiplex Cell Fate Tracking by Flow Cytometry. Methods and Protocols, 3(3):50, July 2020.

[27] Tobias Reichenbach, Mauro Mobilia, and Erwin Frey. Coexistence versus extinction in the stochastic cyclic Lotka-Volterra model. Physical Review E, 74(5):051907, November 2006.

[28] Alexander Dobrinevski and Erwin Frey. Extiction in neutrally stable stochastic Lotka-Volterra models. Physical Review E, 85(5):051903, May 2012.

[29] Samuel Bernard and Hanspeter Herzel. Why Do Cells Cycle with a 24 Hour Period? Genome Informatics, 17(1):72–79, 2009.

[30] S. V. Malinin and V. Y. Chernyak. Transition times in the low-noise limit of stochastic dynamics. Journal of Chemical Physics, 132(1):014504, January 2010.

[31] G. M. Cooper. The Cell: A Molecular Approach. Sunderland (MA): Sinauer Associates, 2 edition, 2000. Chapter: The Eukaryotic Cell Cycle.

